# Decision-related activity and movement selection in primate visual cortex

**DOI:** 10.1101/2023.08.03.551852

**Authors:** Pooya Laamerad, Liu D. Liu, Christopher C. Pack

**Affiliations:** Department of Neurology and Neurosurgery, Montreal Neurological Institute, McGill University, Montreal, Canada

## Abstract

Fluctuations in the activity of sensory neurons often predict perceptual decisions. This connection can be quantified with a metric called choice probability (CP), and there has been a longstanding debate about whether CP reflects a causal influence on decisions, or an echo of decision-making activity elsewhere in the brain. Here we show that CP can actually reflect a third variable, namely the movement used to indicate the decision. In a standard visual motion discrimination task, neurons in the middle temporal (MT) area of the primate visual cortex responded more strongly during trials in which the animals executed a saccade toward their receptive fields, and less strongly for saccades directed away from the receptive fields. The resulting trial-to-trial variability accounted for much of the CP observed across the neuronal population, and it arose through training. Surprisingly, the learned association between MT activity and oculomotor selection was causal, as pharmacological inactivation of MT neurons biased behavioral responses away from the corresponding receptive field locations. These results demonstrate that training on a task with fixed sensorimotor contingencies introduces movement-related activity in sensory brain regions, and that this plasticity can shape the neural circuitry of perceptual decision-making.

## Introduction

In order to make a decision, one must gather information, weigh the evidence in favor of different options, and execute an action. For perceptual decisions, it is tempting to think that each of these steps is supported by different brain regions: Sensory cortices represent the physical properties of stimuli; parietal and frontal cortices accumulate evidence^1^; and motor areas produce actions. While this kind of specialization is to some degree supported by data^2^, the idea of discrete processing stages is probably too simple^3, 4^. Individual neurons in various brain regions contain signals related to multiple aspects of decision-making^5^, and this is particularly so in highly trained animals^6–9^.

For instance, many previous studies have found that, even in sensory cortex^10^, trial-to-trial fluctuations in single-neuron responses predict behavioral responses to identical stimuli^11, 12^. The correlation between these fluctuations and sensory decisions can be quantified with a metric called *choice probability* (CP). Although CP was first reported over 30 years ago, there is still debate about how to interpret it: CP might represent a propagation of random variability from sensory neurons to higher-order structures, or it might reflect feedback about decision that is relayed to the sensory cortex^13, 14^.

Here we consider a third possibility: Fluctuations in sensory neuron activity might be related to *movement selection^a^*, the tendency of neurons to fire more when the subject intends to execute a movement toward their receptive fields^15–17^. Movement selection signals are often found in oculomotor areas, where many cells increase their activity in advance of saccades directed toward particular spatial locations^18^. Such signals are also found in visual cortical areas^19–22^, and like CP, they manifest as fluctuations in single-neuron activity that are independent of the stimulus.

To test this hypothesis, we trained monkeys on a standard visual motion discrimination task and recorded neural activity from the middle temporal (MT) area of visual cortex, which contains neurons that encode motion direction and exhibit CP^11^. We found that CP was closely related to the saccadic eye movement used to indicate the animals’ decisions: The largest CP values occurred when a neuron’s preferred stimulus triggered a saccade toward its receptive field, and CP values were often below chance for saccades directed in the opposite direction. In other words, fluctuations in MT firing conveyed information about future eye movements, and we found that this encoding emerged through training. Moreover, after prolonged training, reversible inactivation of MT led to biases in saccade direction, just as has been found during inactivation of oculomotor brain structures^4, 23, 24^. Our results therefore suggest that CP in simple tasks can reflect movement plans, independent of feedforward or feedback signals related to the stimulus or the decision *per se*. More generally, these results demonstrate how individual brain regions can take on different functional roles through experience.

## Results

We used a variant of a standard discrimination task^11^ to assess the involvement of MT neurons in the perception of visual motion (Figure 1A). Two non-human primates were trained to fixate a small spot at the center of the screen and to view a motion stimulus that was presented in the receptive fields of simultaneously recorded MT neurons. On each trial the stimulus moved in either the preferred or non-preferred direction of the MT neurons, and the animals received a liquid reward for responding to the motion stimulus with a saccade in the corresponding direction (leftward saccade for leftward motion, etc.).

**Figure 1.**
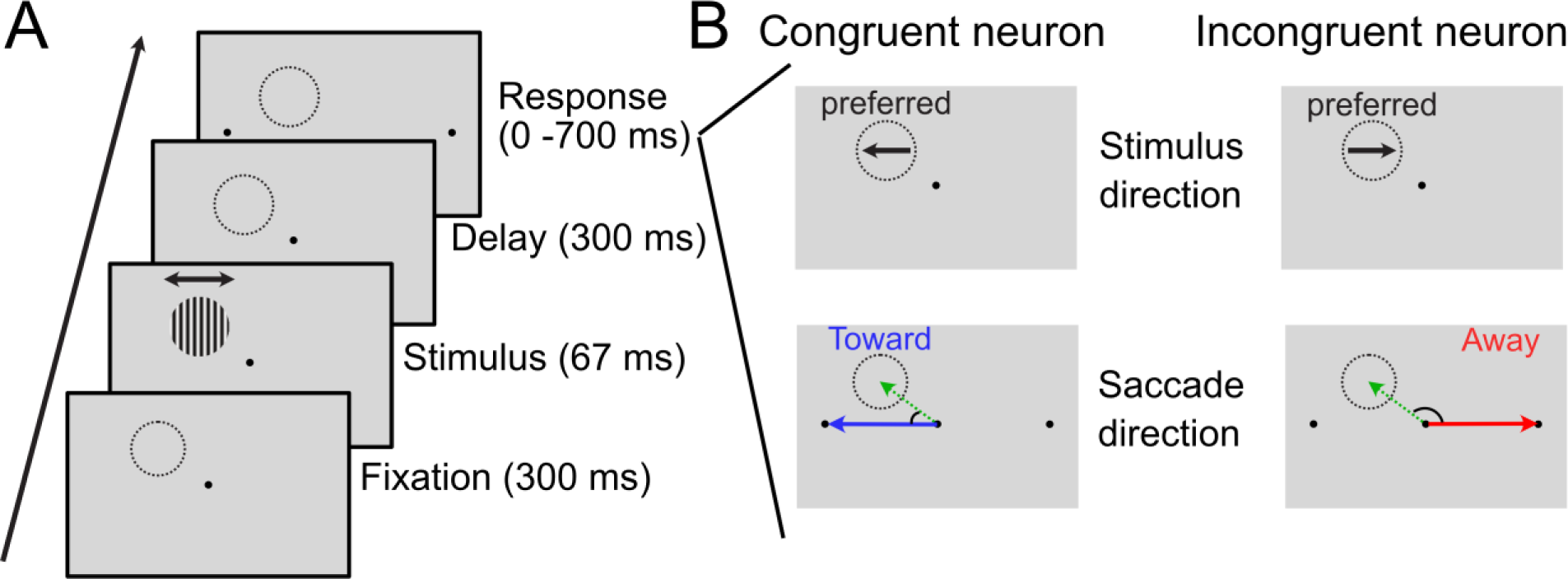
Schematic representation of the experimental paradigm and neuronal categorization. A) Each trial started with a 300 ms fixation period, followed by the presentation of either a drifting grating or random dot stimulus (67 ms) in the receptive field centers. After stimulus presentation, the monkey maintained fixation for an additional 300 ms. Finally, upon disappearance of the fixation point, the monkey made a saccade to one of the two choice targets, indicating its perceived motion direction. B) Categorization of neurons into two groups, congruent and incongruent, based on their preferred directions and the task requirements. Neurons at congruent sites have preferred motion directions that require a saccade towards the approximate center of their receptive field (RF), while neurons at incongruent sites have preferred directions that elicit a saccade away from the RF. The dashed circles represent the RF position; the black arrow indicates the neuron’s preferred motion direction, and the blue and red arrows denote saccade vectors. Green dashed arrow signifies a vector directed towards the RF center.

The animals were trained sequentially to discriminate two different kinds of motion stimuli^25^. In the first phase, the motion stimulus was a Gabor grating, which drifted in either the preferred or anti-preferred direction of the MT neurons. After several weeks of performing this task, the same animals were trained on a second phase, which required direction discrimination of a variable-coherence random dot pattern. After being trained and tested on the dot stimuli, the animals were then retested on the Gabor stimulus. Stimulus duration, size, and speed were identical in both phases, and importantly, both phases of the task imposed a fixed relationship between stimuli and oculomotor responses. At the end of each phase, we performed electrode recordings and reversible inactivation to assess the role of MT in task performance.

### Correlations between MT activity and behavior

CP is a measure of the degree to which an ideal observer could predict behavioral choices based only on the firing rate of a single neuron. For any given stimulus, CP values above 0.5 indicate that firing rates are consistently higher when the subject chooses the response associated with the neuron’s preferred direction, while those below 0.5 indicate higher firing when the subject indicates the non-preferred direction. Most previous studies have reported mean CP values of around 0.55, albeit with considerable variation across neurons^14^.

For continuity with the existing literature, we first present the results for the second phase of our experiment, which made use of the random dot stimuli used in most previous studies. During this phase of the experiment, we recorded single- and multi-unit activity from 129 sites, while monkeys performed the dot motion discrimination task. As in previous studies, CP varied widely across the population, with values ranging from 0.387 to 0.623. The average value was 0.51 ± 0.004, which was significantly greater than 0.5 (p = 0.02, Wilcoxon signed rank (WSR) test).

If CP is related to movement selection, then some of the observed variability should depend on the degree to which the animal intends to saccade toward each neuron’s receptive field. To test this hypothesis, we first categorized recording sites as being either *congruent* or *incongruent*. Congruent sites (n = 64) had preferred motion directions that would trigger a saccade toward the approximate center of their RF (Figure 1B, left). For incongruent sites (n = 65), the preferred motion direction triggered a saccade away from the RF (Figure 1B, right). We split the neurons into these two groups for convenience in illustrating the results, though as shown below, similar conclusions are obtained without any categorization of sites.

Figure 2 shows visual responses from two example neurons, one belonging to the congruent neuronal population (Figure 2A) and the other to the incongruent neuronal population (Figure 2B). Responses are shown when the stimulus motion direction was aligned with the neuron’s preferred direction for both correct (left) and incorrect (right) trials. In both cases, firing rates were higher for stimuli that elicited a saccade toward the RF, even though this was the incorrect answer for the incongruent neuron (bottom right). CP was thus above 0.5 for the congruent neuron and below 0.5 for the incongruent neuron.

**Figure 2.**
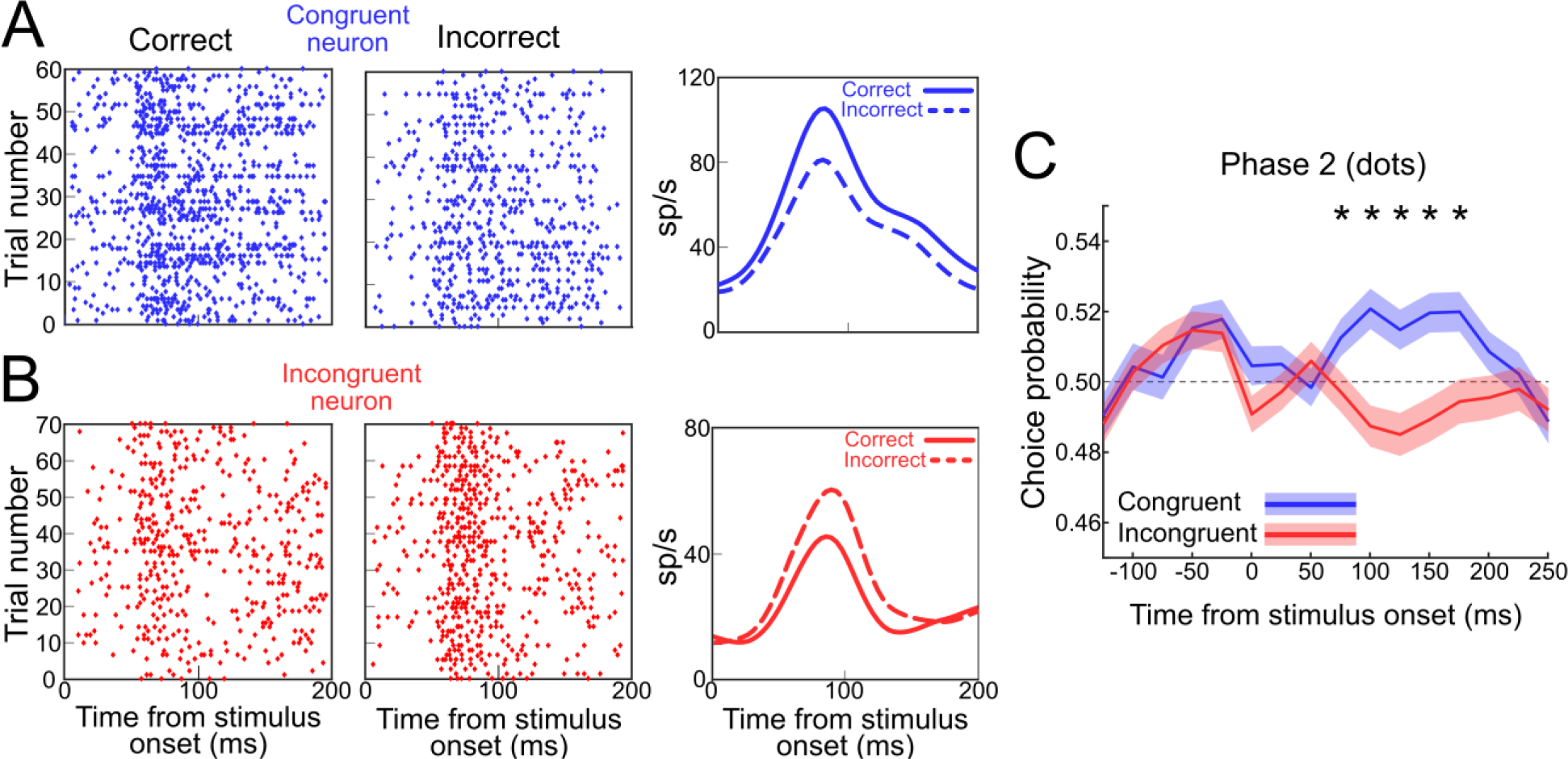
Visual responses of sample neurons and mean of choice probability (CP) for the neuronal population in phase 2 (dots). A) Visual responses of a sample congruent neuron when the stimulus motion direction aligns with the neuron’s preferred direction for both correct (left) and incorrect (middle) trials. The right panel compares the spike density functions of the neuron for correct (solid line) and incorrect (dashed line) trials. B) Example of visual responses from an incongruent neuron. C) Mean CP for all congruent (blue) and incongruent (red) neurons in phase 2 for the random dot stimuls. From 75 ms to 175 ms after stimulus onset, the mean choice probability for the congruent population was significantly greater than that for the incongruent population. Within each panel, responses are aligned on stimulus onset. Shaded colors indicate the standard error of the mean (SEM). Asterisks denote statistically significant differences.

Combining the data across dot coherence levels, the mean CP for the population of congruent sites was 0.527, which was significantly greater than 0.5 (p < 0.01, WSR test), while the mean for the incongruent sites was 0.488, which was significantly lower than 0.5 (p = 0.018, WSR test). The difference in CP between the two populations was highly significant (p < 0.01, Wilcoxon rank sum (WRS) test), even though the informativeness of the neurons about the sensory stimulus (neurometric value) did not differ between the two subpopulations (p > 0.05, WRS test). The results were similar when each animal’s data were considered separately (Supp. Figure 1) and at most individual coherence levels (Supp. Fig. 2; p < 0.01, WRS test).

Figure 2C illustrates the mean CP for the population of congruent (blue) and incongruent (red) neurons, relative to time after the onset of the stimulus. CPs that differed significantly from 0.5 emerged on average with a latency consistent with that of the onset of the sensory response (mean latency ± SEM = 67.69 ± 4.36 ms, Cumulative Sum (CUSUM) analysis), as has been found for visuomotor associations in other brain regions^6, 26^. Thus, they are unlikely to reflect a prior expectation from previous trials, which in any case is not typically found in MT^27^. Nor are they a response to the saccade targets, which did not appear until 100 ms after the time period shown in Figure 2C. Rather, they seem to reflect a learned association between visual stimuli and oculomotor responses. We next examined this hypothesis in more detail.

### Effects of training on choice probability

Because the association between saccade direction and motion direction is imposed by the experimental paradigm, the neuronal effects documented above could have emerged through training. Indeed, training had a strong impact on the role of MT in task performance, as determined through reversible inactivation with the GABA agonist muscimol^25^.

The general effects of inactivation on task performance were described in detail previously^25^. To summarize briefly, after phase 1, reversible inactivation of MT had little effect on task performance^28^. In phase 2, after the animals had spent some weeks training on the random dot stimulus, inactivation of the same MT sites devastated performance on both the random dot and grating task^25^ (see Figure 4 for behavioral results, which will be discussed below). The conclusion of this and related previous work^29, 30^ was that training the animals with random dot stimuli greatly altered the role of MT in perceptual decision-making. The reasons for this are interesting, but not necessarily relevant to this paper (see ^29^ for a detailed discussion). The relevant point here is that the stimulus-related plasticity provides an opportunity to compare CP for the same grating stimuli when MT was minimally involved in perceptual decisions (phase 1) and when it was strongly involved (phase 2).

We recorded neural activity in response to grating stimuli from 75 sites in phase 1 and 81 sites in phase 2. In contrast to previous work on perceptual learning^8^, we did not find that average CP values increased with training (phase 1: mean ± SEM = 0.511 ± 0.006; phase 2: mean ± SEM = 0.513 ± 0.006; p > 0.05, WRS test). This is not necessarily surprising, because unlike in previous studies, overall performance on the grating task did not change between phase 1 and phase 2^8^.

However, when the data were analyzed with respect to the distinction between the congruent and incongruent MT populations, a clear effect of training on CP emerged. Figure 3 compares the mean time course of CP for congruent neurons (blue; n = 36) and incongruent neurons (red; n = 39), for phase 1 (panel A) and phase 2 (panel B) of the experiment. For phase 1, there was no significant difference between the average CP of the two populations at any time point (p > 0.05, WRS test). In contrast, during phase 2 of the experiment, the mean CP was above 0.5 for congruent neurons (blue; n = 35) and below 0.5 for incongruent neurons (red; n = 46) at time points from 75 ms to 175 ms after stimulus onset (p < 0.01, WRS test). This was true in each animal individually (Supp. Fig. 3) and for most individual stimulus contrast levels (Supp. Fig. 4). These differences were not due to changes in the experimental configuration across phases, as there were no significant differences in the RF positions (p > 0.05, WRS test) or preferred directions of the neurons (p > 0.05, WRS test) recorded in the two phases of the experiment. We also did not detect any change in the sensitivity of MT neurons to the grating stimuli between phases^25^.

**Figure 3.**
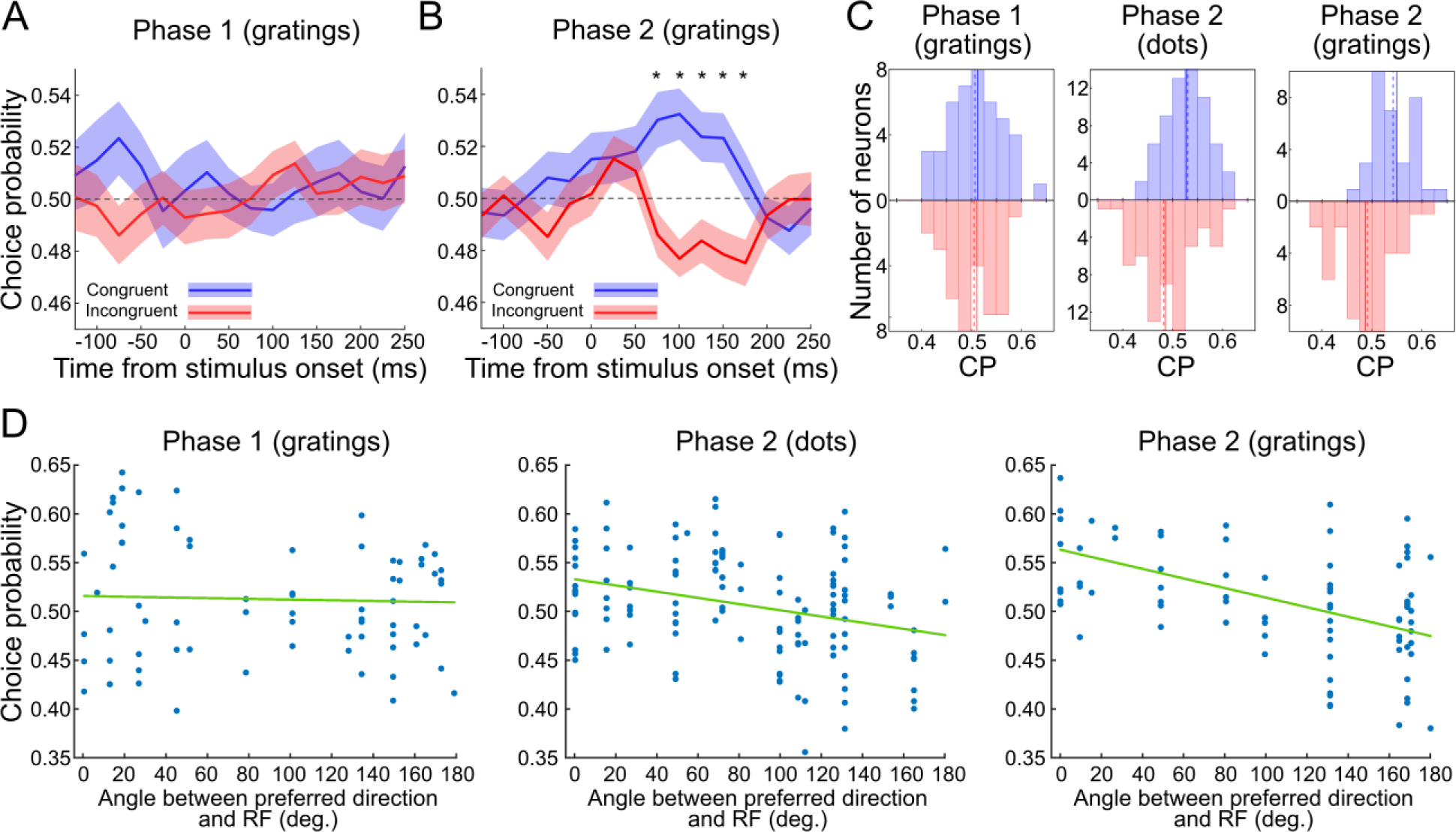
Comparison of mean CP for congruent and incongruent neurons across experiment phases (grating). A) During phase 1 (grating), there was no significant differences between levels of CP in the congruent and incongruent populations. B) Testing with gratings during phase 2 revealed a significant difference in CP between the two populations at time points ranging from 75 ms to 175 ms post-stimulus. C) CP for congruent and incongruent populations across different phases of training: For phase 1, the mean CP of congruent neurons (blue, mean CP ± SEM = 0.513 ± 0.009) and incongruent neurons (red, mean CP ± SEM = 0.512 ± 0.007) are not significantly different (mean difference = 0.001, p > 0.05, WRS test); 2) For phase 2 with dot stimuli, the mean CP for the congruent population (blue, mean CP ± SEM = 0.527 ± 0.005) was significantly higher (mean difference = 0.039, p < 0.01, WRS test) than for the incongruent population (red, mean CP ± SEM = 0.488 ± 0.006); 3) For gratings in phase 2, the difference between the mean CP of congruent (blue, mean CP ± SEM = 0.55 ± 0.008) and incongruent (blue, mean CP ± SEM = 0.487 ± 0.007) populations was also significant (mean difference = 0.063, p < 0.01, WRS test). D) CP plotted against the angle between each neuron’s preferred motion direction and a vector connecting the fovea to the RF center. In phase 1, there was no significant correlation between CP and this angular difference (correlation coefficient: −0.03, p > 0.05). However, phase 2 showed a significant correlation, for both dot and grating stimuli, indicating larger CPs for neurons whose preferred motion direction aligned more closely with their RF position (correlation coefficient for dot: −0.30, p < 0.01; for grating: −0.50, p < 0.01). Shaded colors indicate the standard error of the mean (SEM). Asterisks denote statistically significant differences. Solid lines represent the mean of the data, while dashed lines indicate the median values.

Averaging the neural data across a time window of 75 – 175 ms after stimulus onset provides a summary of the distribution of CP values (Figure 3C) for the two phases of the experiment and both stimulus types. Statistically, there was a significant interaction between cell type and experimental phase (ANOVA; F (1,304) = 7.11, p < 0.01). Importantly, this relationship between time-averaged CP and congruency remains if congruency is defined along a continuum, rather than categorically. To visualize this relationship, we plotted CP against each recording site’s degree of *congruency* (Fig. 3D), defined as the angle between the receptive field position and the preferred visual motion direction (Figure 1B). This revealed a strong, negative correlation during phase 2 of the experiment for both dots (r = −0.30, p < 0.01) and gratings (r = −0.50, p < 0.01). During phase 1, the relationship was also negative but far weaker, and not statistically significant by our criteria (correlation coefficient: 0.04, p > 0.05). Similar correlations and training effects were observed when congruency was defined as the Euclidean distance between the RF position and the nearest saccade target (Supp. Fig. 5). The relationship between CP and neurometric sensitivity did not appear to vary with training, for either the congruent or incongruent populations (p > 0.05, Fisher’s F test; Supp. Fig. 6).

### Causal relationship between MT and behavioral biases

These results suggest that CP is shaped by training and that it is at least partially related to movement selection. As with decisions, CP in MT could reflect a causal influence on movement selection, or it could indicate that the results of the selection process are fed back to MT. If the influence is causal, then inactivation of MT should bias behavioral responses in such a way as to favor saccade responses directed away from the inactivated RFs, irrespective of the direction of stimulus motion. Such effects are often found in oculomotor areas like the superior colliculus, lateral intraparietal area, and the frontal eye fields^23, 24, 31–34^.

We examined the effect of muscimol inactivation of the same populations of MT neurons described above. As described previously, muscimol effectively abolished action potentials near the site of injection, with peak behavioral effects occurring roughly 18 hours later^25, 35^. At this time point, muscimol had spread over roughly 2 mm of cortex^35^, affecting a retinotopic zone covering the RFs in our recordings.

During phase 2 of the experiment, inactivation of MT led to a clear behavioral bias toward saccade targets positioned opposite the inactivated RFs, for both gratings (Figure 4A) and dots (Supp. Fig. 7A) (p < 0.05, WRS test). This bias was not present before inactivation in phase 2 or during inactivation in phase 1 (Figure 4B; H (3) = 18.81, p = 0.031, Kruskal-Wallis test, FDR corrected). These effects of inactivation were found in each animal individually (Supp. Fig. 7B) and evident across most levels of grating contrast (Supp. Fig. 7C). As reported previously, they were accompanied by large increases in behavioral thresholds for motion discrimination^25^, indicating that MT in phase 2 was involved in both motion discrimination and movement selection.

**Figure 4.**
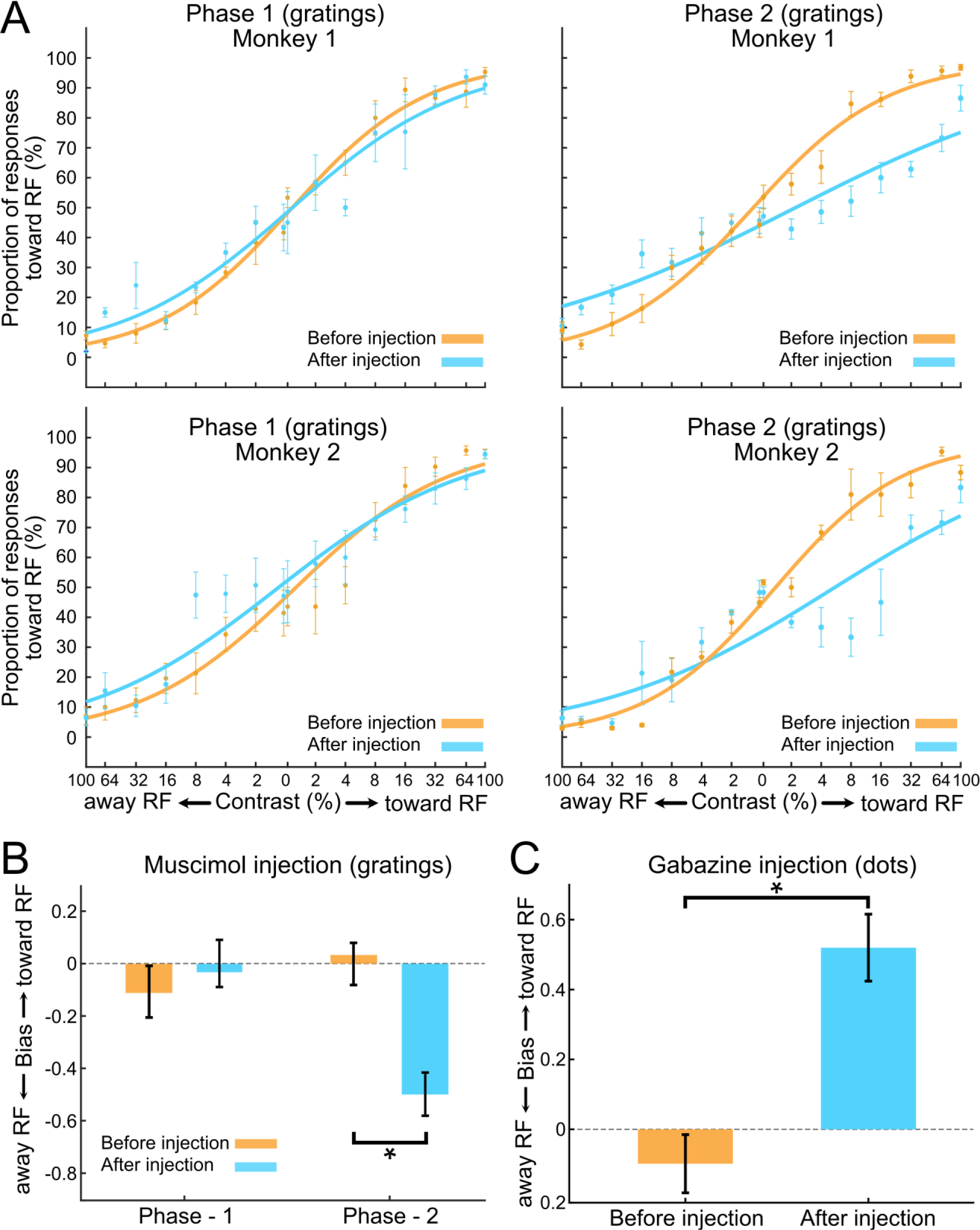
Training effects on behavior. A) Following the training of phase 2, monkeys developed a bias to make saccades away from the inactivated receptive field (RF). During phase 1, MT inactivation did not lead to consistent biases toward or away from the RFs. During phase 2, muscimol injections led to a clear bias for saccades away from the RF. B) In phase 2, injection of muscimol led to a significant bias away from the RFs (H (3) = 18.81, p = 0.031, Kruskal-Wallis test, FDR corrected) Positive values in this plot represent a bias toward the RF and negative values represent a bias away from the RF. C) In phase 2 for dot stimuli, after injection of gabazine, the animals developed a bias to make saccades toward the RFs of the injected sites (p < 0.01, WRS test). Error bars indicate the standard SEM. Asterisks denote statistically significant differences.

In one animal, we carried out a third phase of training that exclusively involved Gabor gratings (120 sessions). After this training period, the behavioral bias revealed by muscimol injections decreased significantly (p = 0.021, WRS test; Supp. Fig. 7D) and was no longer statistically different from zero. Thus, the role of MT in movement selection depended heavily on the nature of the training, rather than its overall duration, just as reported previously for discrimination performance^25, 35^.

Overall, these results suggest that pharmacologically-induced decreases in MT firing rates (to zero) were associated with a propensity to saccade to targets away from the inactivated RFs. Thus, the neural results reported above likely reflect a causal influence of MT activity on movement selection. To examine this issue further, we performed additional experiments in which we artificially *increased* the firing rate of MT neurons, by injecting the GABA antagonist gabazine into the MT sites^36^. As expected from a causal influence, this led to a behavioral bias *toward* the corresponding RFs, for dot stimuli in phase 2 (Figure 4C; p < 0.01, WRS test). This effect of gabazine injection was found in each animal individually (Supp. Fig. 7E; p < 0.01, WRS test).

### Further considerations on the effects of inactivation

It is possible that these behavioral biases reflected an influence of inactivation on the visibility of the corresponding saccade target, rather than a process of movement selection. We consider this explanation to be unlikely because the visibility of the target was the same in phase 1 and phase 2, which yielded very different effects of inactivation. Moreover, in phase 2 the bias was largely absent for the lowest Gabor stimulus contrasts (Supp. Fig. 7C; p > 0.05, WSR test), which would not be expected if it was caused by decreased visibility. Previous work has found that MT lesions do not impair the ability to make saccades to stationary targets^37^.

These results also cannot be interpreted in terms of a bias to report particular motion directions, as our data set contained roughly equal numbers of congruent and incongruent sites. Nor can they be explained by a general impairment in the ability to understand or to perform the task. As noted previously^25^, the effects of inactivation were specific to the trained stimulus location and to the retinotopic site of the muscimol injection. Moreover, during inactivation, the number of “failed” trials, in which the animals did not saccade to either target, did not differ when the correct response was toward vs. away from the RF (p > 0.05, WSR test; Supp. Fig. 8). Thus, the results shown in Figure 4 are more consistent with biased movement selection than with a general impairment in task performance.

### Stimulus coding

As noted above, training did not lead to appreciable changes in neural sensitivity or readout efficiency. However, previous work has found that training is associated with large changes in the trial-to-trial variability of neural responses^38^. Because such variability is necessary for detection of CP^39^, we wondered whether it also changed with training in our experiment.

Indeed, within the population of congruent neurons, there was a significant increase in variability, as quantified by the Fano factor, from phase 1 to phase 2 of the experiment (p < 0.01, WRS test; Figure 5A). This increased variability was shared among congruent neurons, leading to a large increase in pairwise noise correlations within this population (p = 0.011, WRS test; Figure 5B). Similar changes were not found in the incongruent population, and were not detectable when the MT population was considered as a whole^8^ (p > 0.05, WRS test). The lack of changes in the incongruent population is somewhat unexpected, although as noted in the Discussion, it may reflect constraints imposed by the anatomical projections from oculomotor areas to visual cortex^40^.

**Figure 5.**
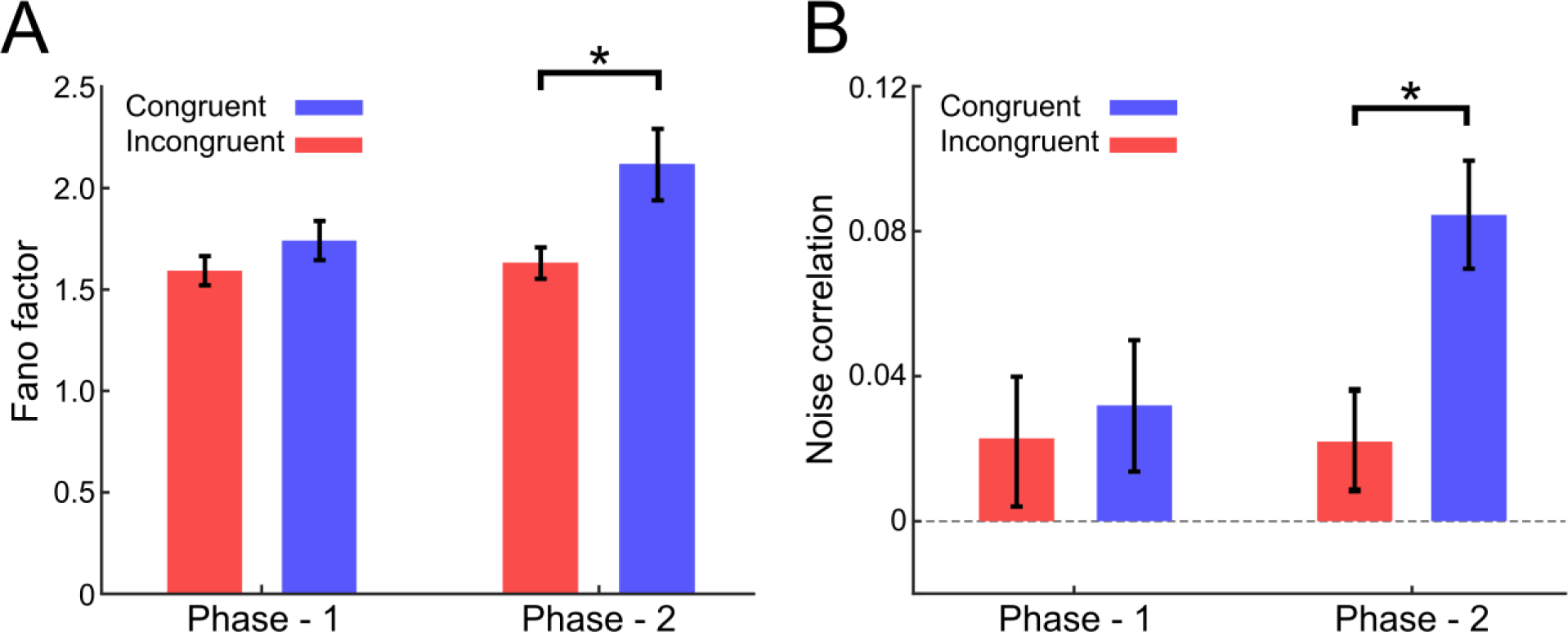
A) Fano factor of congruent neurons increased with training. In phase 1, there was no significant difference between the Fano factors of congruent and incongruent populations. However, in phase 2, a significant increase was seen in the Fano factor for the congruent population, and this increase was significantly different from that of the incongruent population (p = 0.011, WRS test). B) In phase 2, there was a significant increase in the noise correlations within the congruent population, and it was significantly larger compared to those observed in the incongruent population (p < 0.01, WRS test).

## Discussion

We have shown that CP in a standard motion discrimination task is closely related to movement selection, a well-known modulatory influence on visual responses. CP values were higher for neurons whose receptive fields were near the future target of a saccade, and often below 0.5 for neurons whose receptive fields were opposite the direction of an impending saccade (Figure 2). This relationship between CP and movement selection emerged with training (Figure 3) and coincided roughly with the appearance of a causal influence of MT neurons on movement selection (Figure 4A). Put simply, after appropriate training, a portion of the firing rate in MT seemed to encode movement direction, independently of the visual stimulus. This led to increased firing rate variability, which was shared across a subpopulation of MT neurons (Figure 5). Because many behavioral tasks make use of an eye movement as the perceptual readout, we speculate that similar effects are present in some previous studies of the relationship between neural activity and perceptual decisions.

### Comparison with previous work

CP was initially thought to reflect a causal influence of sensory signals on perception^11^, and indeed it is often still interpreted this way. In this conception, random variation in the neural response to a stimulus is transmitted directly or indirectly to higher-order brain regions, biasing decisions accordingly. An alternative idea is that CP reflects an echo of a perceptual decision that is made elsewhere in the brain and fed back to sensory cortex^10, 12, 13^. The precise nature of choice-related activity has been shown with reversible inactivation to be highly task-dependent^33, 41^, at least in parietal cortex, but the situation in sensory cortex is less clear^42–44^. Recent computational analyses suggest that both feedforward and feedback influences might be found at the same time in the same sensory neurons^45, 46^.

Our results suggest that there is a component of CP that is neither feedforward nor feedback, in the conventional (functional) sense of the words. That is, signals related to movement selection neither reflect strictly the properties of the sensory input nor the decision made by the subject. The latter distinction is clearest in incongruent sites in MT (Figure 2), which fire more strongly when the animal chooses a response that is inconsistent with their preferred visual stimulus.

It is difficult to say how much this third factor might have affected previous reports on MT, as RF positions are seldom reported. On the one hand, most studies report mean values of CP in MT that are above 0.5, independently of any consideration of RF positions^8, 11, 47^, which suggests that there is likely a component of CP that is unrelated to movement selection. This is evident in our data as well (Figure 3). Moreover, several studies have found significant CP in the absence of a visual stimulus or a saccade^48^, and in the absence of a consistent spatial relationship between stimulus and response^49^. Thus, it seems unlikely that movement selection accounts entirely for CP in all previous studies.

On the other hand, the existing literature describes substantial variations in CP levels across tasks and even across individuals performing the same task^11, 50, 51^. Our results suggest that one source of such variability is the locations of the RFs relative to the movements demanded by the task. For area MT, these factors might be inherently correlated, as there is a small bias in direction preferences toward centrifugal motion (i.e., motion directed away from the fovea), even in untrained animals^52^. As centrifugal motion is encoded by congruent cells in most motion tasks, this bias could produce above-chance CP values, even in the absence of activity that is truly decision-related. Similar correlations between receptive field position and stimulus preferences are sometimes found in V1^53^, the ventral visual pathway^54^ and the subcortex^55^. Some experiments that did not impose a straightforward relationship between stimuli and motor responses have yielded little or no CP^51, 56, 57^.

One limitation of our study is that we used an unusually brief motion stimulus. This might explain why our mean CP was at the low end of published values for motion discrimination tasks in MT^50^, though it must be noted that longer stimulus durations can inflate CP values, via microsaccades^58^. More importantly, the short stimulus duration limits our ability to analyze the dynamics of CP signals. Previous work with brief motion pulses revealed responses in parietal cortex that were consistent with a feedforward model of evidence accumulation^59, 60^. But studies that varied the timing of motion pulses within a longer stimulus duration often found that CP was more consistent with feedback, as its magnitude increased through time^12, 45, 60^. Note that this latter observation is also consistent with a role for movement-related signals, which similarly increase through time within a trial^61^. In any case, we suspect that movement selection effects are present in some published results on CP (see ^62^ for a likely example), and it would be interesting to examine this factor in existing data, where possible.

### Nature of movement selection signals

Many perceptual discrimination n paradigms could be considered variants of the antisaccade task that is commonly used in oculomotor studies: a peripheral stimulus triggers a saccade either toward or away from the receptive field^26^. Neuronal responses in these tasks often contain a mix of visual and task-related signals^63–67^, the latter often occurring with latencies comparable to those of the feedforward visual response^68^, just as we have reported here (Figure 2C).

For these kinds of tasks, a likely source of movement-related signals is the superior colliculus (SC)^18^, which has indirect, reciprocal connections with the visual cortex and plays an important role in target selection^34^. The retinotopic nature of these projections^40^ would explain why congruent neurons seem to be specifically affected by training (Figure 5): The SC neurons that project to MT encode saccades into the contralateral visual field, which is only true of congruent neurons in our task. Similarly, training induces selectivity primarily for “congruent” visual motion in the superior colliculus^7^, along with other oculomotor areas^6, 39, 69^, indicating a bidirectional connection between sensory and motor areas that is shaped by training but limited by anatomy.

Despite this connection, the movement selection signals we have observed are probably not strongly related to the movements themselves. Even in highly trained animals, microstimulation of MT can alter the endpoint of a saccade^70^, but it does not by itself trigger a saccade^71^. Thus, the modulatory effects we have observed are more likely due to the linkage between stimulus and response than to the saccadic response itself^33^.

This kind of modulation is often associated with covert spatial attention. Attentional influences could in principle explain why, in our data, congruent cells exhibit an increase in firing rate when the saccade target is near the RF and a decrease when it is far away (Figure 2)^72^. It could also explain why tasks that dissociate the correct motor response from the stimulus yield CP that is spatially unselective^49^. At the same time, it is difficult to explain the below-chance CP we have observed for incongruent neurons in this way. Shifts of attention take hundreds of milliseconds to manifest in cortex^73^, which is presumably too long to account for our results (Figure 3). The fact that noise correlations and trial-to-trial variability increase with CP modulation (Figure 5) also argue against an explanation of our results in terms of spatial attention^74, 75^.

### Implications for neural coding

Some previous studies of sensory plasticity have used perceptual learning paradigms, wherein training leads to improvements on the corresponding behavioral task. Such learning usually leads to relatively modest changes in neural tuning or sensitivity, even when behavioral performance improves more substantially^8, 76 77, 78^. Changes in noise correlations also occur with training^38^ and task^42^, though they tend not to be structured in such a way as to explain changes in behavior^14, 38, 79^.

Our experiments did not involve perceptual learning, as there was no detectable change in performance on the Gabor grating task between phase 1 and 2 of the experiment^25^. Thus, we interpret the increase in noise correlations that we observed for congruent MT neurons (Figure 5) in terms of the introduction of a new kind of input to visual cortical neurons, in this case movement-related. This kind of input is not likely to improve sensory encoding^80^ or decision-making^10^.

The fact that the changes in noise correlations were only detectable in the pool of congruent neurons points to a somewhat unusual interpretation of the relationship between learning and decision-making. It is possible that the animals in our study treated the task as one of detecting certain kinds of motion signals, rather than discriminating between them. Specifically, they might have simply attempted to detect motion directed away from the fovea, which would necessarily entail monitoring the congruent neurons. Although this might seem like an odd strategy, the representation of motion in area MT is biased toward such stimuli^52^, as is human perception^81^. Previous work has shown that subjects often solve discrimination tasks by exploiting or creating biased representations^82, 83^.

More generally, these results highlight the role of experimental design choices in shaping the resulting neural data. A task like ours, which can be solved by a simple mapping of stimuli to movements, probably encourages the formation of a rather unnatural, low-dimensional decision space^82^. Thus, a better understanding of the neural circuitry for decision-making might require more naturalistic paradigms that contain a richer variety of stimuli and responses^84, 85^.

## Methods

### Experimental setup

All experimental methods were described in our previous papers^25, 86^; most of this paper entails a reanalysis of the corresponding data. Briefly, monkeys were implanted with devices for head fixation and electrode recordings, then trained to perform a standard motion discrimination task (Figure 1). Recordings were performed with a linear electrode array (Plexon V-Probe) that was positioned so as to record from area MT, which was identified based on anatomical MRI scans, electrode depth, and the prevalence of strong, direction-selective visual responses. Eye movements were tracked at 1000 Hz with an infrared tracker (EyeLink).

In some sessions, muscimol (typically 2 μL at 0.05 μL/min; concentration of 10 mg/mL) was injected through a fluid channel in the V-Probe, to inactivate nearby neural activity. As described in our previous paper^25^, we conducted various controls to verify that muscimol silenced neural activity, that the effects were localized to the injection site, that they were caused by muscimol and not by the injection *per se*, and that the procedures and inactivation effects did not vary across experimental phases^25^. For gabazine (8 sessions in monkey 1 and 6 sessions in monkey 2), we used injections of ∼0.5 μl at 0.05 μl/min. and a concentration of 0.05 mm.

### Behavioral paradigm and visual stimuli

The experiment consisted of two phases. In the first phase, animals were trained to report the motion direction of a drifting grating. Once behavioral thresholds had stabilized, we performed 10 sessions (4 sessions in monkey 1 and 6 sessions in monkey 2) of neuronal recording and pharmacological inactivation of area MT. In the second phase, the same animals were trained to perform the same task, but with random dot kinematograms as the stimuli. Again, neuronal recordings and inactivation were performed after behavioral training was complete. During this phase, animals were also tested with drifting gratings for comparison with the first phase. Training in each phase lasted 3 – 4 weeks. One animal performed a third phase of training with only gratings (Supp. Figure 7D), which lasted several months.

Stimulus parameters, such as the spatial and temporal frequency of the Gabor and the speed of the dots, were chosen to match the preference of the MT neurons near the recording site. The motion direction was always chosen according to the preferred or anti-preferred direction of the neurons under study. The size of the stimulus also matched the RF size of the MT neurons (mean radius = 6.3° ± 1.2°). The contrast of the Gabor patch or the coherence of the random dots pattern was chosen randomly on each trial from among seven or eight values spanning the range of the monkey’s psychophysical threshold. These values were selected based on previous measurements of the psychometric functions with an emphasis around the steepest sections of the psychometric function. For the grating stimuli, these values did not change across experimental phases, as performance remained stable after the initial training in phase 1.

On each trial, animals acquire fixation for 300 ms, after which the motion stimulus was presented briefly (typically for 67 ms). The monkeys were then required to maintain fixation for another 300 ms, after which the fixation point disappeared, two choice targets appeared, and the monkey made a saccade to the corresponding target to report its perceived motion direction (preferred or null relative to the neurons isolated; Figure 1). Targets were typically positioned at 10° eccentricity, with angles sampled at 45° intervals. The correct saccade direction was linked to the most similar saccade target direction (i.e. rightward saccade for rightward motion, etc.). The monkey was required to indicate its decision within 700 ms following the onset of the choice targets. Correct choices were rewarded with a drop of liquid. For the trials that contained no motion signal (0% contrast or 0% coherence), rewards were delivered randomly on half the trials. If fixation was broken at any time during the stimulus, the trial was aborted. In a typical experiment, the monkeys performed the task on 20-40 repetitions of each stimulus.

### Data analysis

#### Neural data

We considered both single-neuron and multi-unit activity, as CP does not generally differ for these types of signals^43, 44, 79, 87^. Choice probability was computed using standard methods^11, 88^, in which neuronal firing rates were z-scored for each stimulus condition and time bin. We then formed two distributions of firing rates corresponding to the animal’s behavioral responses in the task and subjected these data to ROC analysis. Choice probability was then calculated as the area under the ROC curve. CP values were combined across stimulus conditions (typically 2, 4, 8% contrast or 0, 2, 4, 8% coherence) for which there were a sufficient number of incorrect behavioral responses to distinguish behavioral choice from stimulus-driven responses.

Neurons were characterized as congruent or incongruent based on the angle between their preferred motion direction and a vector connecting the fixation point to the receptive field center. If the absolute value of this angle was less than 90°, the cell was considered congruent; otherwise, it was incongruent.

To analyze the dynamics of choice-related activity (Figures 2 and 3), we first computed the CP for each neuron across stimulus conditions, spanning from 300 ms before to 300 ms after stimulus onset, using a window size of 50 ms and a step size of 5 ms. The emergence of statistically significant CP was detected with the Cumulative Sum (CUSUM) algorithm^89^.

Firing rate variability was estimated using spike counts taken from a 100-ms window, spanning the time period between 75 ms and 175 ms following stimulus onset. The Fano factor, calculated as the ratio of spike-count variance to spike-count mean, was determined using the Variance Toolbox ^90^. Data for each neuron at every coherence level were analyzed independently, after which the variance (across trials) and the mean of the spike count were calculated.

Noise correlations were calculated as the Pearson correlation coefficient representing the trial-by-trial covariation of responses from pair of neurons that were recorded simultaneously^80^. Each neuron’s responses were z-scored by subtracting the average response and dividing by the standard deviation across multiple stimulus presentations. This procedure eliminated the impact of stimulus strength and direction on the average response, enabling noise correlation to solely capture correlated trial-to-trial fluctuations around the mean response. To avoid correlations influenced by extreme values, only trials with responses within ±3 standard deviations of the mean were considered^80^.

#### Behavioral data

Behavioral thresholds were measured as the contrast (for gratings) or coherence (for dots) required to obtain 82% correct performance, based on Weibull fits to the corresponding psychometric functions. To evaluate bias, we compared the monkeys’ performance when the motion stimulus triggered a saccade towards the receptive field (RF) versus when it drove a saccade away from the RF, using the congruent and incongruent definitions introduced above. We separately calculated the proportion of correct performance (%) for both saccade directions (toward and away from RF) and applied the equation below to determine bias^91^:

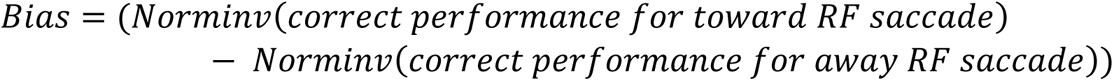

Positive values indicate a bias to make saccades toward the RF, while negative values indicate a bias away from the RF.

### Statistics

We applied nonparametric analyses for statistical evaluation of our data, with the exception of determining significant interactions between experimental phases and types of neurons. For this specific computation, we employed a two-way ANOVA, as it is the most suitable method for assessing significance of interactions. Following this, whenever multiple comparison across conditions were required for any analysis, we used the Tukey’s Honestly Significant Difference method to adjust for multiple comparisons by maintaining the false discovery rate (FDR) at 5% for all tests.

## Acknowledgements

This work was supported by a grant from the CIHR (PJT178071) to CCP. We thank Dr. Ralf Haefner for helpful discussions.

## Supplementary figures

**Figure S1.**
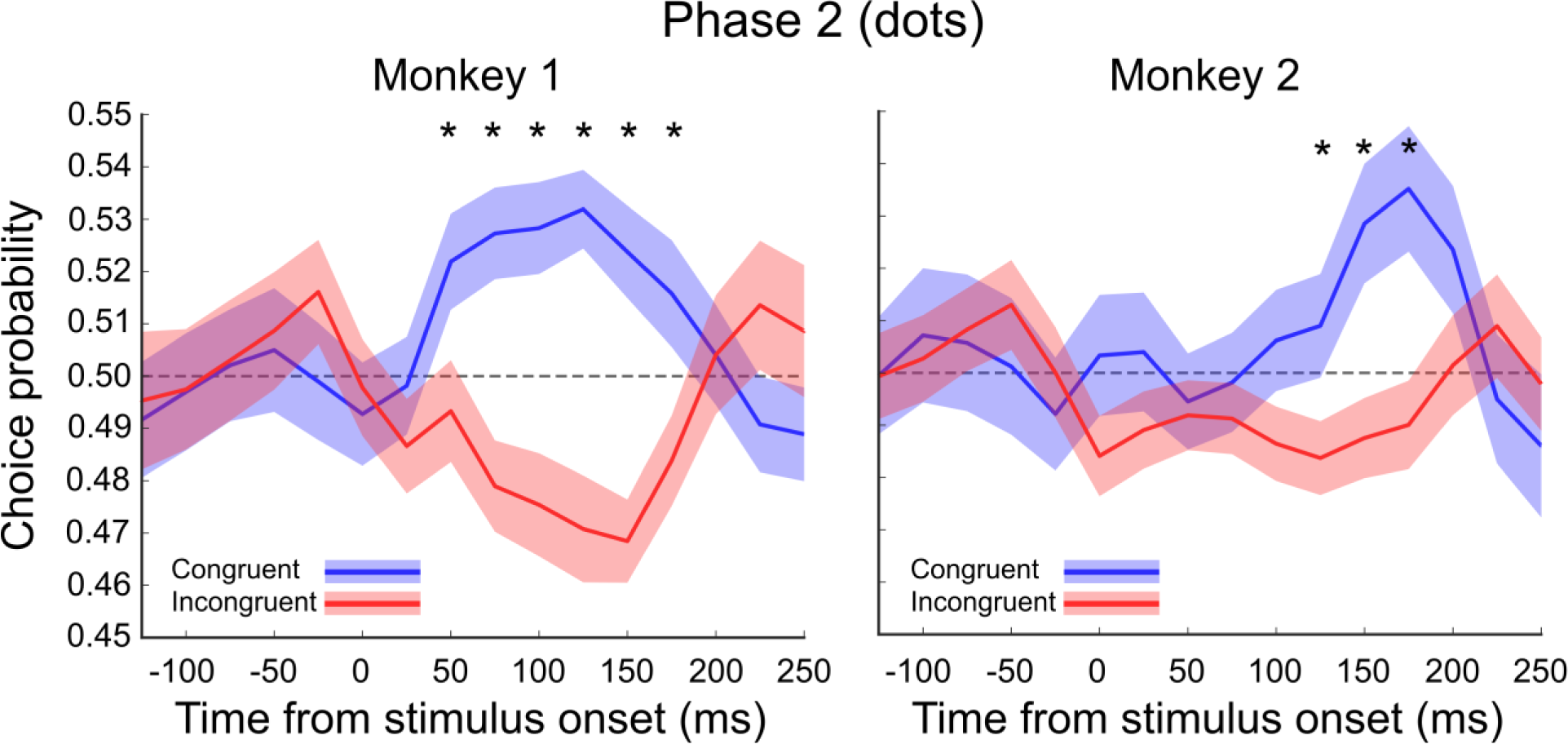
Comparison of mean choice probability (CP) for congruent and incongruent neurons in each monkey during the second phase (dot). CP values were calculated individually for each monkey, with blue representing congruent populations and red indicating incongruent populations. The mean CP significantly differed in Monkey 1 (left) between 50 and 175 ms after stimulus onset (p < 0.05; Wilcoxon Rank-Sum test), while in Monkey 2 (right), the difference was significant between 100 and 175 ms after stimulus onset (p < 0.05; Wilcoxon Rank-Sum test). Shaded colors indicate the standard SEM. Asterisks denote statistically significant differences.

**Figure S2.**
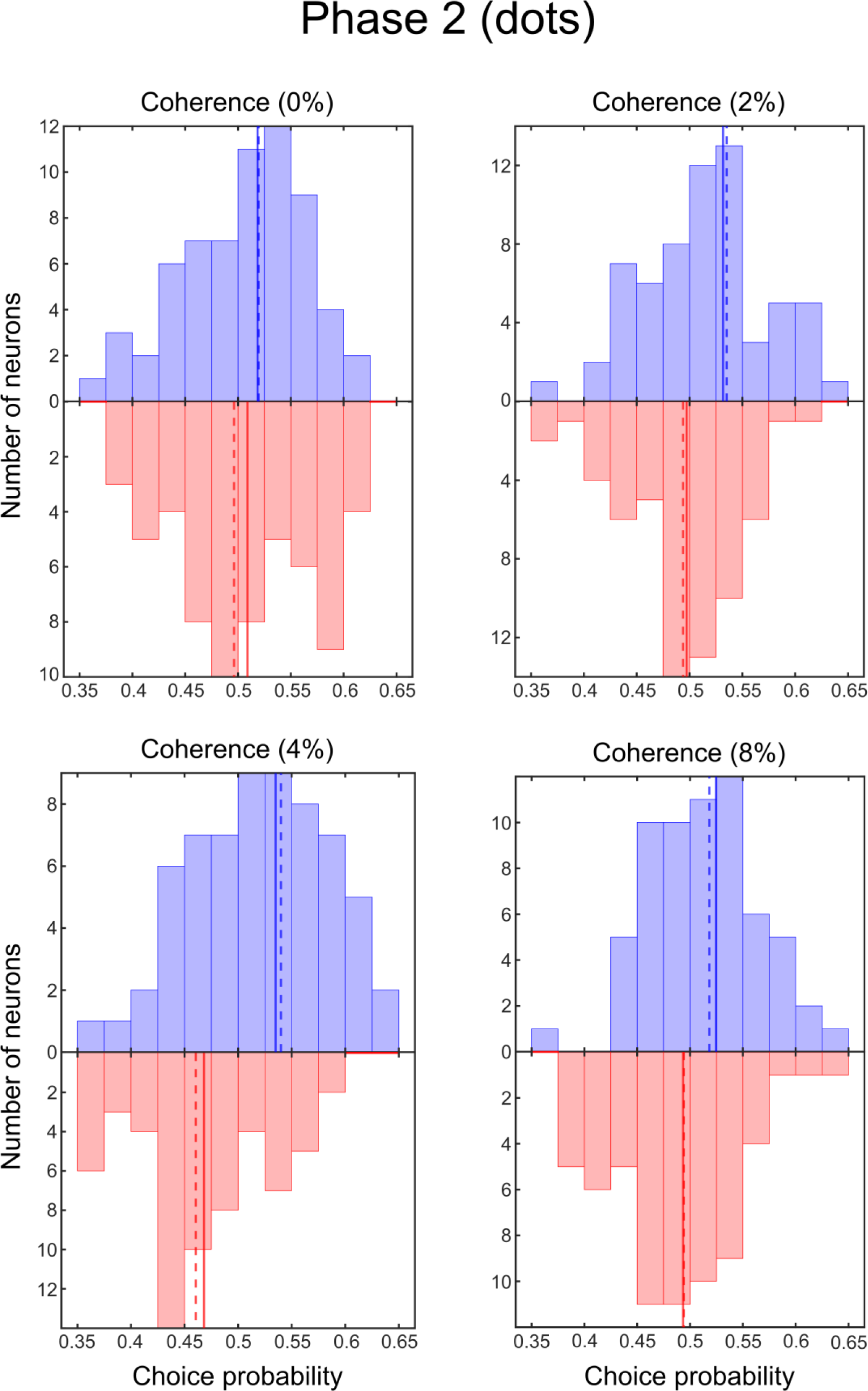
Distribution of choice probability (CP) values in each coherence in the second phase (dot). CP values were calculated for congruent (blue bars) and incongruent (red bars) neuronal populations by averaging neural data over a 75-175 ms time window following stimulus onset and computing CP for each coherence level individually. The median CP for the congruent population was higher than that of the incongruent population in all coherence levels, with the difference being significant for all levels except 0% coherence. (Coherence 0%: congruent mean = 0.522, incongruent mean = 0.511, p > 0.05; coherence 2%: congruent mean = 0.532, incongruent mean = 0.492, p = 0.016; coherence 4%: congruent mean = 0.537, incongruent mean = 0.467, p < 0.01; coherence 8%: congruent mean = 0.525, incongruent mean = 0.494, p = 0.048; Wilcoxon Rank-Sum test). Solid lines represent the mean of the data, while dashed lines indicate the median values.

**Figure S3.**
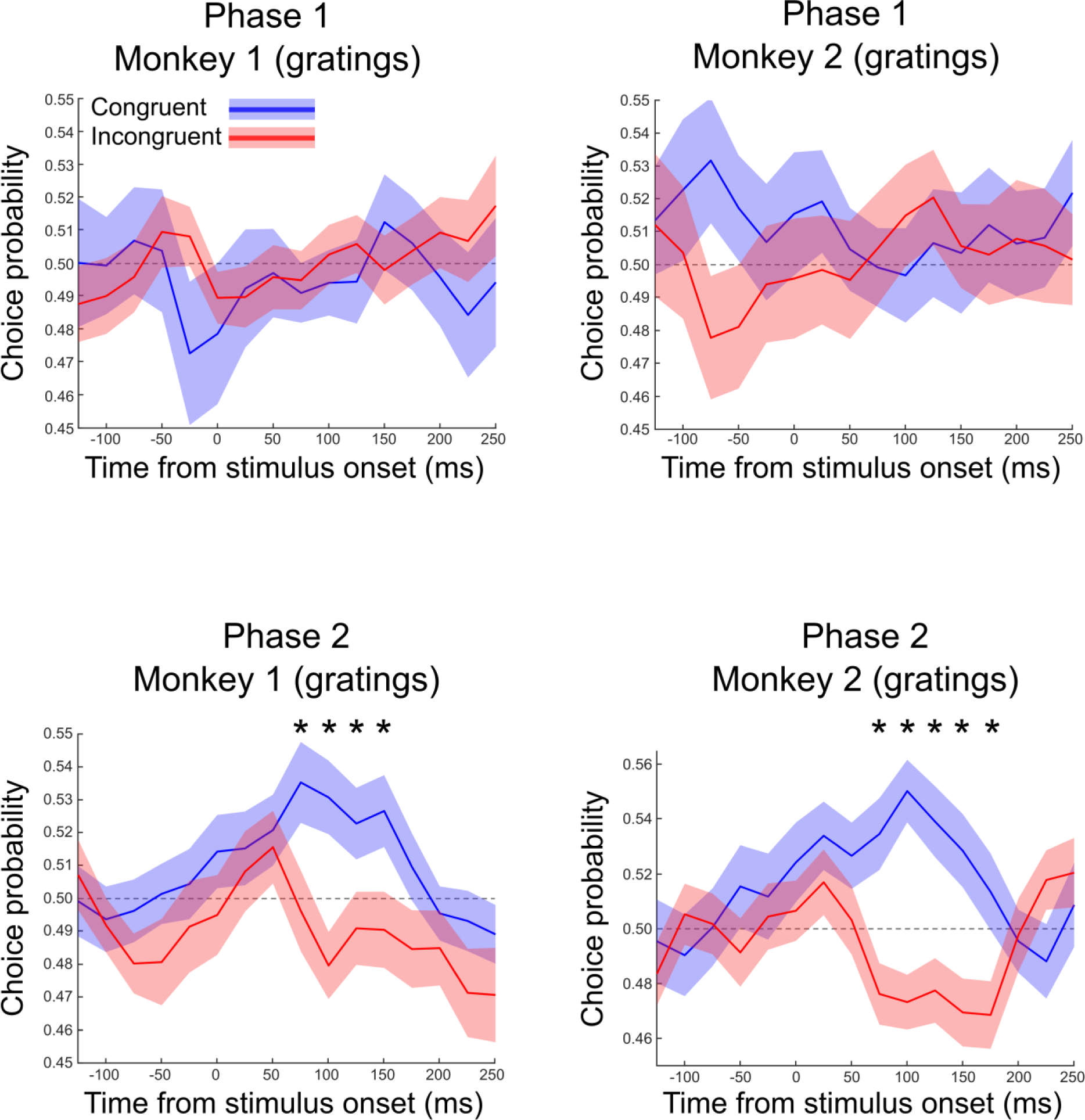
Comparison of mean choice probability (CP) for congruent and incongruent neurons in each monkey across two phases (grating). We computed CP values individually for each monkey, with congruent populations denoted by blue and incongruent populations by red. During phase 1, no significant difference was observed between the two populations, as shown in top tow of the figure. However, in phase 2, a significant difference in the mean CP was found in Monkey 1 between 75 and 150 ms post stimulus onset, as shown in bottom row of the figure. (p < 0.05; Wilcoxon Rank-Sum test). For Monkey 2, a significant difference was found between 75 and 175 ms following the stimulus onset (p < 0.05; Wilcoxon Rank-Sum test). The shaded regions represent the standard SEM. Asterisks denote statistically significant differences.

**Figure S4.**
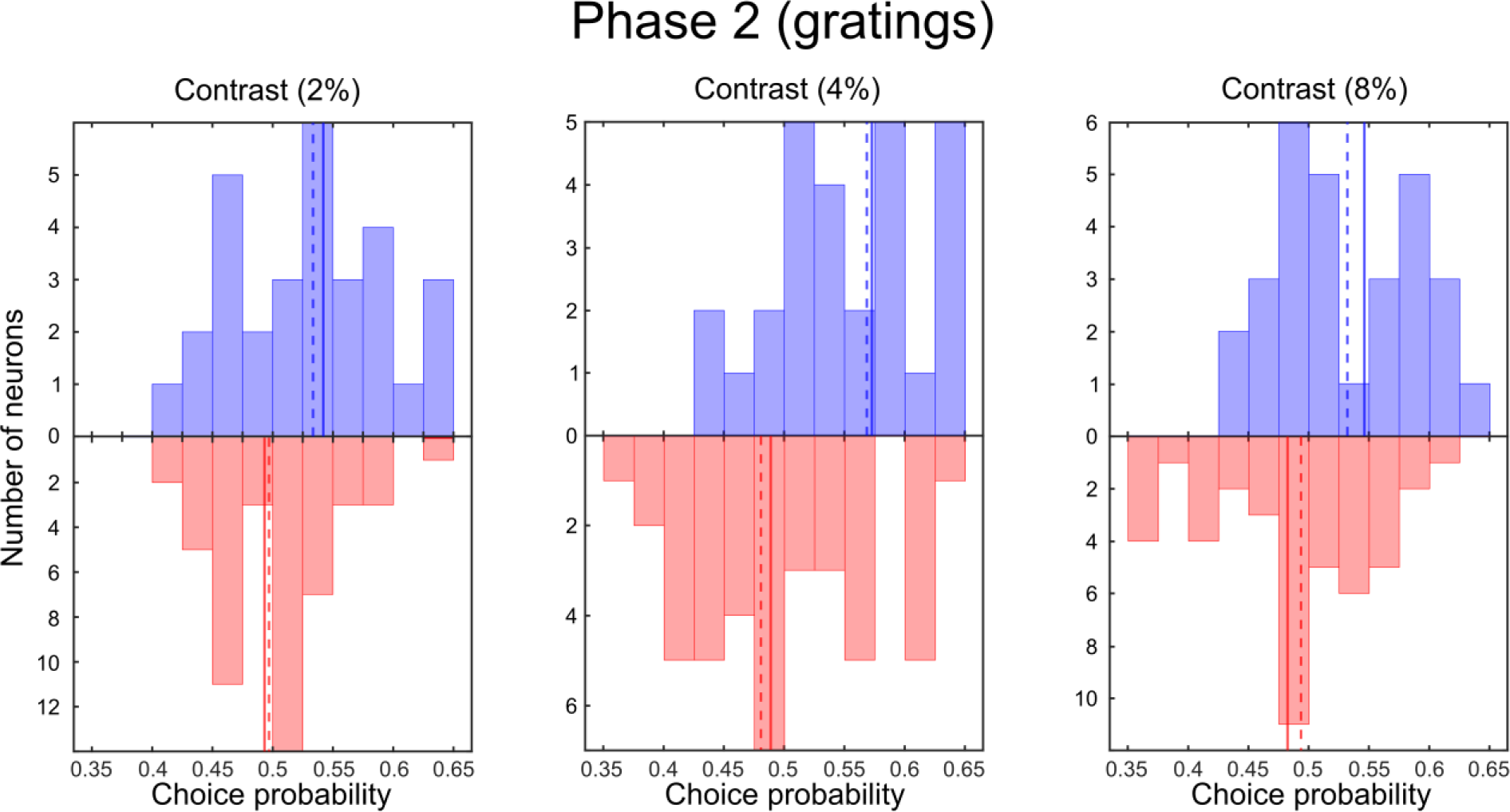
Distribution of choice probability (CP) values in each coherence in the second phase (grating). CP values were calculated for congruent (blue bars) and incongruent (red bars) neuronal populations by averaging neural data over a 75-175 ms time window following stimulus onset and computing CP for each contrast level individually. The mean CP for the congruent population was higher than that of the incongruent population in all contrast levels, with the difference being significant for all levels (contrast 2%: congruent mean = 0.54, incongruent mean = 0.49, p < 0.05; contrast 4%: congruent mean = 0.57, incongruent mean = 0.49, p < 0.01; contrast 8%: congruent mean = 0.54, incongruent mean = 0.48, p < 0.05; Wilcoxon Rank-Sum test). Solid lines represent the mean of the data, while dashed lines indicate the median values.

**Figure S5.**
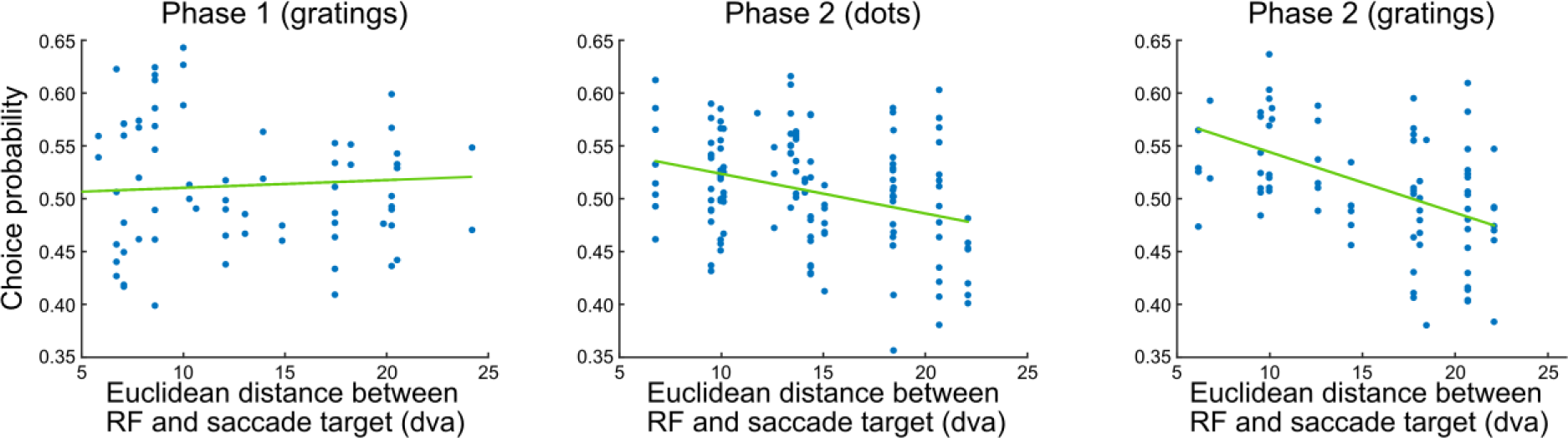
Analysis of the neurons’ choice probability (CP) based on the Euclidean distance between the estimated RF center and the saccade target. During phase 1, no significant correlation was observed between the neurons’ CP and their distance (correlation coefficient: 0.05, p > 0.05). In contrast, phase 2 revealed a significant correlation between CP and distance. The correlation was stronger for grating stimuli compared to random dots (correlation coefficient for dot: −0.31, p < 0.01; correlation coefficient for grating: −0.48, p < 0.01).

**Figure S6.**
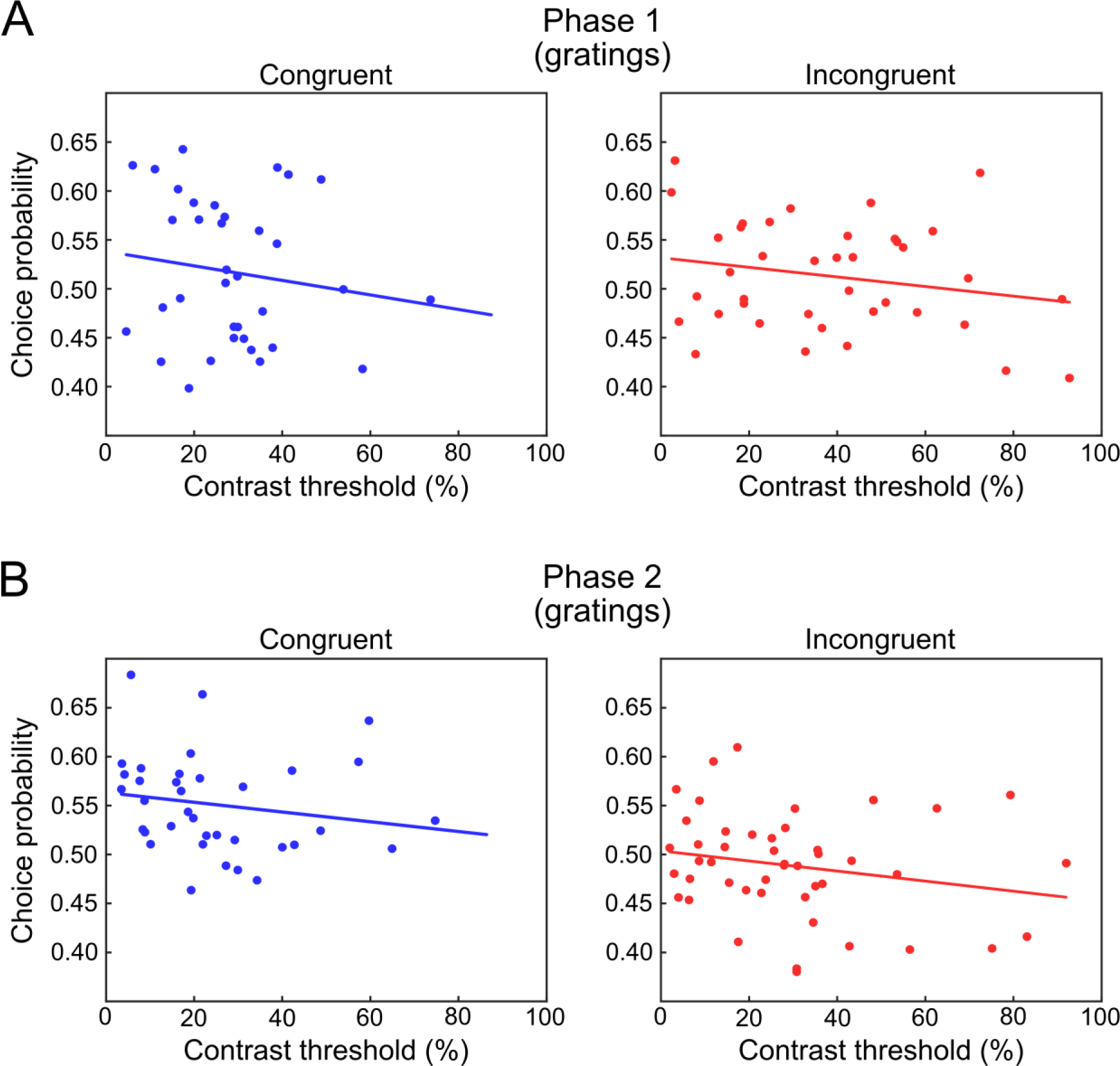
Relationship between CP and neurometric thresholds for grating stimuli in each subpopulation and in the two different experimental phases. A) In phase 1, there was no significant difference in the slopes of the regression between CP and neurometric threshold for congruent (blue) and incongruent (red) neurons (congruent: r = −0.16, p = 0.31; incongruent: r = - 0.20, p = 0.22, Fisher’s test for correlation difference: z = 0.17, p > 0.05). B) Similarly, slopes were also not significantly different between the two populations (congruent: r = −0.17, p = 0.33; incongruent: r = −0.21, p = 0.15, Fisher’s test for correlation difference: z = 0.09, p > 0.05). The intercepts did differ between the two populations in phase 2 (z-value = 3.17, p < 0.01, z-test) but not in phase 1 (z-value = 0.21, p > 0.05, z-test)

**Figure S7.**
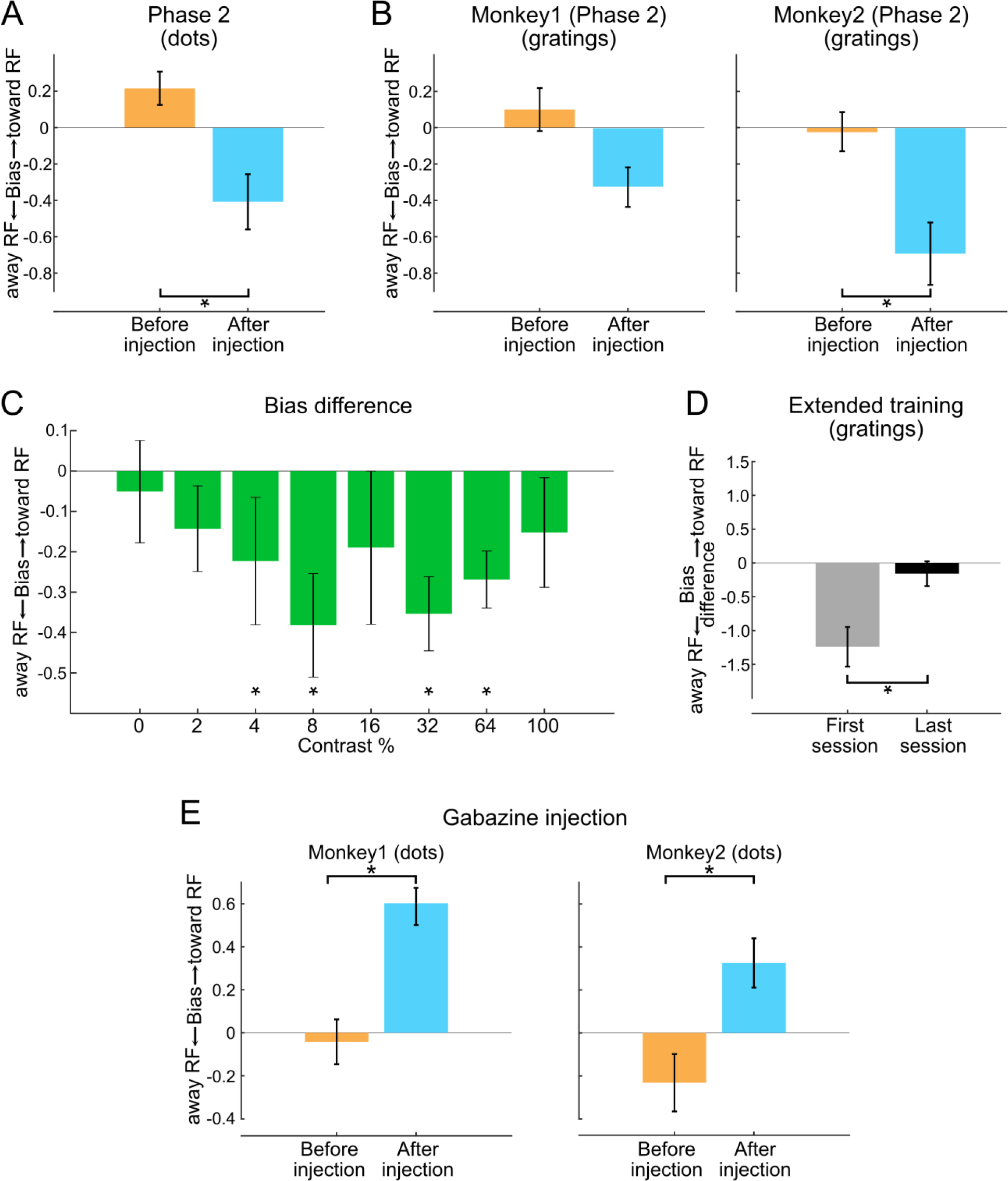
Evaluating saccade bias before and after muscimol and gabazine injections. A) In phase 2, muscimol injections led to a bias for saccades away from the RF for dot stimuli (p = 0.036, WRS test). B) This shift in saccade bias after muscimol injection was found in each monkey individually for grating stimuli. In both monkeys, the difference of bias before and after inactivation was significant (Monkey 1: p = 0.032, Monkey 2: p < 0.01, WRS test). C) We computed the difference in saccade bias before and after muscimol injection across all contrast levels during phase 2 for grating stimuli. For contrast levels of 4%, 8%, 32% and 64%, the difference in bias was significantly different from zero (p < 0.05, WSR test). D) An additional 120 sessions of training of the second monkey with grating stimuli led to a reduction in the difference in saccade bias before and after muscimol injection. This difference was significantly smaller in the final session compared to the first (p = 0.021, WRS test). (E) The injection of gabazine led to a significant shift in the monkeys’ bias to make saccades toward the RF. This shift in saccade bias after gabazine injection was found in each monkey individually for dot stimuli. In both monkeys, the difference of bias between before and after inactivation was significant (Monkey 1: p < 0.01, Monkey 2: p < 0.01, WRS test).

**Figure S8.**
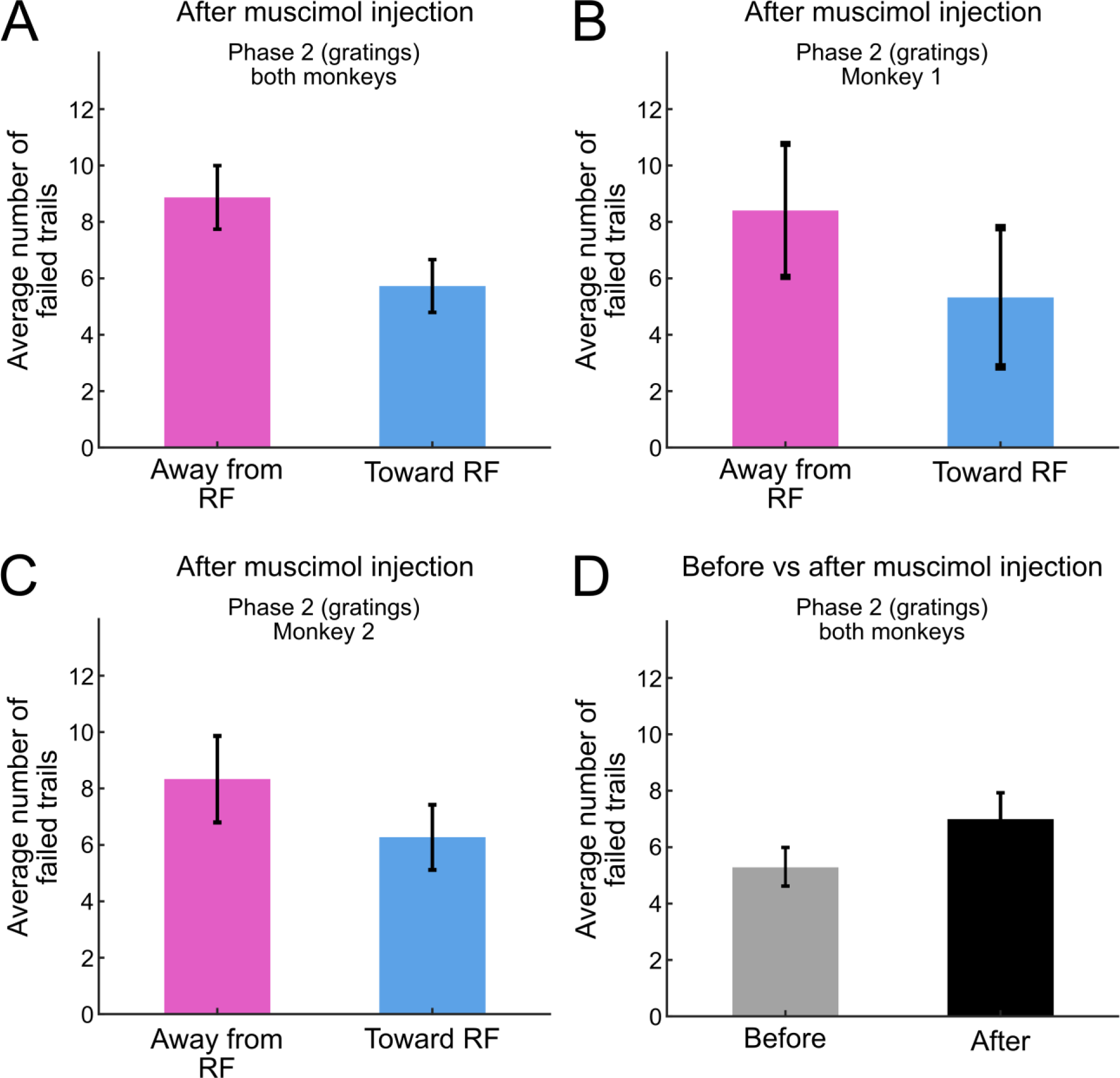
Average number of failed trials. A) In each session following the muscimol injection during phase 2 for grating stimuli, we considered the number of trials in which the monkey successfully maintained fixation and the stimulus was presented, yet the monkey failed to successfully complete the trial by selecting a target. We refer to these as “failed trials”. Comparing the number of failed trials when the correct response was towards the RF versus when it was away from the RF in each session, at each coherence level, revealed no significant difference in the number of failed trials (P > 0.05, WSR test). B and C) An individual analysis in each monkey also showed no significant difference in the number of failed trials (P > 0.05, WSR test). D) A comparison of the number of failed trials before and after muscimol injection sessions for grating stimuli in phase 2 showed a slight increase in the average number of failed trials after the injection, but the difference was not statistically significant (P > 0.05, WSR test).

## Footnotes

a We use the term “movement selection” somewhat arbitrarily, as our experiments were not designed to distinguish between selection of spatial locations, targets, and saccades (see Sato & Schall, 2003).

